# SiGra: Single-cell spatial elucidation through image-augmented graph transformer

**DOI:** 10.1101/2022.08.18.504464

**Authors:** Ziyang Tang, Tonglin Zhang, Baijian Yang, Jing Su, Qianqian Song

**Author notes:** Corresponding authors: Qianqian Song, PhD; Jing Su, PhD; Baijian Yang, PhD.

## Abstract

The recent advances in high-throughput molecular imaging push the spatial transcriptomics technologies to the subcellular resolution, which breaks the limitations of both single-cell RNA-seq and array-based spatial profiling. The latest released single-cell spatial transcriptomics data from NanoString CosMx and MERSCOPE platforms contains multi-channel immunohistochemistry images with rich information of cell types, functions, and morphologies of cellular compartments. In this work, we developed a novel method, Single-cell spatial elucidation through image-augmented Graph transformer (SiGra), to reveal spatial domains and enhance the substantially sparse and noisy transcriptomics data. SiGra applies hybrid graph transformers over a spatial graph that comprises high-content images and gene expressions of individual cells. SiGra outperformed state-of-the-art methods on both single-cell spatial profiles and spot-level spatial transcriptomics data from complex tissues. The inclusion of immunohistochemistry images improved the model performance by 37% (95%CI: 27% – 50%). SiGra improves the characterization of intratumor heterogeneity and intercellular communications in human lung cancer samples, meanwhile recovers the known microscopic anatomy in both human brain and mouse liver tissues. Overall, SiGra effectively integrates different spatial modality data to gain deep insights into the spatial cellular ecosystems.

## INTRODUCTION

Recent advances in spatial molecular imaging have allowed examining the spatial landscapes and transcriptional profiles of complex tissues at subcellular resolution^1-3^. The interrogation of the spatial locations and gene expressions of individual cells within a tissue aid in understanding spatial heterogeneity of cell-to-cell communications and cell interactions with its surrounding environment, which is crucial for understanding disease pathology. Current commercially available technologies for single cell spatial profiling, such as NanoString CosMx™ Spatial Molecular Imager (SMI)^4^ and the Vizgen MERSCOPE/MERFISH platforms^5,6^, are capable to accurately capture the locations of targeted transcripts, cell locations, and cell boundaries, accompanied with multi-channel immunohistochemistry (IHC) images. For example, the NanoString CosMx™ is capable to simultaneously assay up to 1,000 genes^4^ and 100k to 600k cells per slide, dramatically exceeding current single cell omics technologies. Therefore, the emerging single-cell spatial transcriptomics (SCST) commercial platforms and are revolutionizing current spatial biology research, holding the promises to spatially and functionally reveal complex architectures within tissues, and furthering our insights into the underlying disease mechanisms at unprecedented resolution^7-9^.

The emerging SCST multi-modal data provides new opportunities for accurately identifying spatial domains, which is crucial for revealing and functional annotating the cellular anatomy of complex tissues. Existing methods for deciphering spatial cell clusters still rely on the clustering methods for non-spatial single-cell RNA-seq data, such as the Seurat^10^ and the Louvain clustering based Scanpy^11^ method that only take gene expression data as input. Other methods have been developed to include spatial information to improve the identification of spatial regions. For example, stLearn^12^ leverages gene expression of neighboring spots and tissue image features to identify the spatially distributed clusters. BayesSpace^13^ enables spatial clustering through a Bayesian statistical method with the joint analyses of gene expression matrix and spatial neighborhood information. SpaGCN^14^ identifies spatial regions using graph convolutional network, with the spatial graph constructed from gene expression and histology information. Though these methods show their capability in spatial clustering, the power of different modalities within single-molecule spatial imaging profiles is not fully unleashed to achieve desirable performance.

In addition to domain recognition, the enhancement of spatial gene expression data also presents a significant challenge. Though great progress has been made in spatial technologies, the major problems such as missing values, data sparsity, low coverage, and noises^2,15^ encountered in spatial transcriptomics profiles are impeding the effective use and the elucidation of biology insights^16,17^. Meanwhile, the multi-channel spatial images in single-cell spatial data consist of high-resolution, high-content features detected in the tissue, such as cell types, functions, and morphologies of cellular compartments, as well as the spatial distributions of cells. Incorporating such imaging features into transcriptomics data processing will help address the challenges of missing values and alleviating expression noise, thus enhance the spatial transcriptomics data quality for downstream analytical tasks.

In this study, we developed a novel SiGra method, i.e., SIngle-cell spatial elucidation through image-augmented GRAph transformer, to decipher spatial domains and enhance spatial signals simultaneously. SiGra is one of the first method to utilize multi-modalities including multi-channel images of cell morphology and functions to address technology limitations and achieve augmented spatial profiles. SiGra accurately recovers missing information in spatial gene expressions, uncover cellular dynamics, and reveal spatial architecture of cellular heterogeneity within tissues. Through the extensive and quantitative benchmarking with existing methods on multiple datasets including both single-cell level and spot-level spatial data generated by different platforms, SiGra demonstrates superior performance in terms of spatial domain identification, latent embedding, and data denoising. Collectively, SiGra will contribute to uncovering the complex spatial architecture within heterogeneous tissues and facilitate the gaining of biological insights. SiGra is provided as an open-source software available at https://github.com/QSong-github/SiGra, with detailed tutorials demonstrating the applications on different SCST platforms.

## RESULTS

### Overview of the SiGra method

The SiGra method includes 1) the graph representation of the original spatial transcriptomics data (**Fig. 1a**), and 2) the hybrid graph transformer model to elucidate the spatial patterns and enhances the raw gene expression data (**Fig. 1b**).

**Fig. 1:**
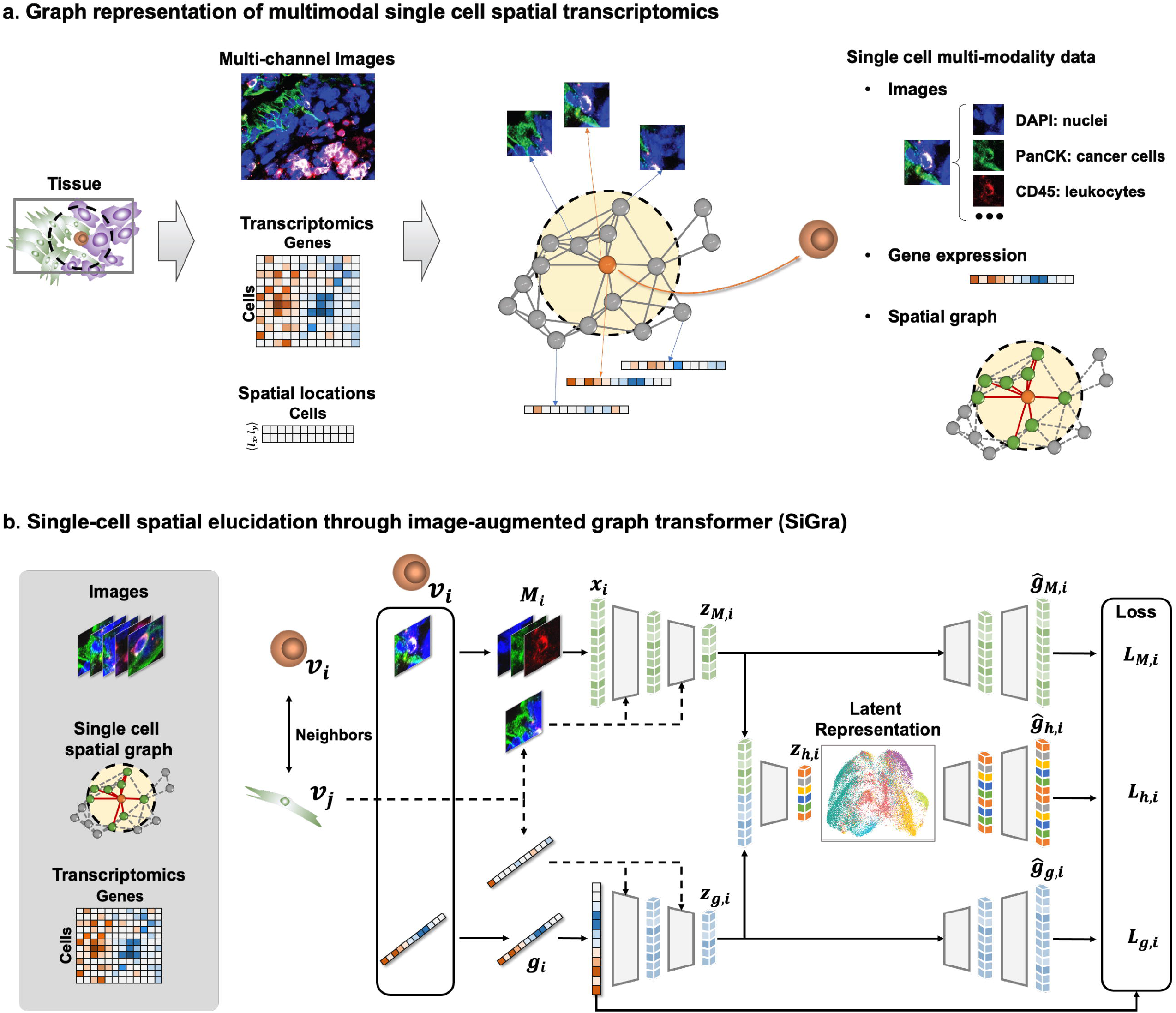
Schematic overview of the SiGra method. **a**, Graph representation of the spatial transcriptomics profiles. Each cell on the constructed spatial graph is accompanied with its multi-channel images and gene expression. **b**, SiGra comprises three graph transformer autoencoders (imaging, transcriptomics, and hybrid, respectively) with attention mechanism to incorporate the single-cell multi-modal data for simultaneous data enhancement and spatial domain recognition.

### Graph representation

The state-of-art SCST data consists of 1) the multi-channel images of biomarkers for cell types (e.g., pan-cytokeratin or PanCK staining for tumor cells, CD3 for T cells, and CD45 for leucocytes) and cell compartments (e.g., DAPI staining for cell nuclei and CD298 staining for cell membrane). For each staining channel, a high-content grayscale image is assembled from a series of Field of View (FOV) images; 2) the vendor-provided cell segmentation results such as the coordinates of cell centroids and the hull of cell boundaries; and 3) the cell-level summarization of gene expression according to the coordinates of each detected transcript and the cell boundary identified from cell segmentation.

In SiGra, the single-cell spatial graph is constructed based on the spatial centroid of detected cells, with each node representing a cell, and each edge representing two neighboring cells (Euclidian distance shorter than 14 – 16μm). Each node/cell within the spatial graph is accompanied with multi-modal data (images and gene expression) extracted from the original spatial profiles. Specifically, for each cell, an image of 21.6-by-21.6μm centered at the cell centroid is cropped from each immunohistochemistry (IHC) image. For example, as NanoString CosMx data consists of five channels (DAPI, PanCK, CD45, CD3, and CD298), each cell is associated with five single-cell images. In this way, SiGra achieves the graph representation of spatial profiles, i.e., the single-cell spatial graph with each located cell’s multi-channel images and gene expression.

### Hybrid graph transformer model

The SiGra model comprises three graph transformer autoencoders (imaging, transcriptomics, and hybrid, respectively) with attention mechanism (**Fig. 1b**) to incorporate the single-cell multi-modal data for simultaneous data enhancement and spatial domain recognition. Regarding the imaging autoencoder, with a cell *i* represented by node *v*_*i*_, an array of single-cell IHC images *M*_*i*_ is converted to a vector ***x***_i_, and projected to the latent space as ***Z***_*M, i*_ through multi-head graph transformer layers (**Supplementary Fig. 1, Materials and Methods**). This latent imaging features ***Z***_*M,i*_ then reconstructs the gene expression profile 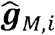 of cell *i*. For the transcriptomics autoencoder, the same architecture is used for the the latent imaging features representation (*Z*_*g,i*_) and the reconstruction 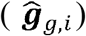 of the original expression ***g***_*i*_ in cell *i*. For the hybrid autoencoder, the latent imaging features (***Z***_*M,i*_) and the latent expression features (***Z***_*g,i*_) are concatenated and projected as a hybrid feature ***Z*** _*h,i*_, which is used to reconstruct the gene epression 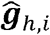 of cell *i*. Imaging and gene expression features of neighboring cells, represented as neighbor nodes *v*_*j*_ ∈ 𝒩 (*v*_*i*_) in in the spatial graph, are also used as the input for graph transformers, so that the spatial cellular information is aggregated into the model.

SiGra learns the reconstructed gene expression via a self-supervised loss of combined MSE from gene embedding *L*_*M,i*_, image embedding *L*_*g,i*_ and combined embedding *L*_*h,i*_, with the loss function

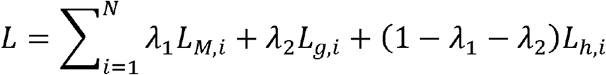

where the hyperparameters *λ*_1_, *λ*_2_ ≥ *λ*_1_ + *λ*_2_ ≤ 1 and *N* is the total cell number. After training, SiGra outputs the hybrid reconstruction 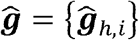 as the final enhanced expression profile. The latent representation. ***Z=*** {***Z***_*h,i*_}of the original SCST data is used for spatial data clustering.

With the introduced multi-head attention mechanism in graph transformer layers, SiGra adaptively updates the contributions of neighboring cells {*v*_*j*_} to cell *v*_*i*_ through aggregating and propagating the extracted image features and the gene expression features from neighbors, eventually updates the latent representation of cells and the final reconstructed gene expression profiles. Through the evaluation and benchmarking with current available methods, SiGra demonstrates exceptional performance on multiple spatial transcriptomics datasets from different platforms, especially on single-cell spatial profiling. Moreover, the enhanced spatial transcriptomics data by SiGra facilitates the insights into cellular communications and the underlying biological discoveries.

### SiGra accurately identifies spatial domains in the single-cell spatial profiles of NanoString CosMx SMI

To evaluate the performance of SiGra in deciphering spatial domains, we compare it with five state-of-the-art clustering methods developed specifically for spatial transcriptomics, including Seurat v4^10^, Scanpy^11^, stLearn^12^, SpaGCN^14^, and BayesSpace^13^. For comparisons, we use the SCST dataset of Lung-9-1 generated by NanoString CosMx SMI. This dataset consists of 20 Field of Views (FOVs) from lung cancer patient tumor tissue^4^, with 982 genes and 83,621 cells that covers majorly eight cell types, including lymphocytes, neutrophile, mast, endothelial, fibroblast, epithelial, myeloid, and tumors. Details of these experimental data are provided in Data Availability. The identified spatial clusters of each method are annotated based on the matched overlap of spatial clusters and ground truth manually annotated in the original study^4^.

**Fig. 2a** shows the ARI scores of all the 20 FOVs. Notably, SiGra is shown to identify the most accurate spatial clusters (median ARI: 0.59) than the rest of methods, especially higher than stLearn (median ARI: 0.22) and Scanpy (mean ARI: 0.25). Compared with SpaGCN (mean ARI: 0.27), Seurat (median ARI: 0.38) and BayesSpace (mean ARI: 0.32) present relatively better performance with identified clusters more consistent with manual annotations. These comparison results demonstrate that SiGra improves the identification of spatial clusters than existing methods for single-cells spatial profiles.

**Fig. 2:**
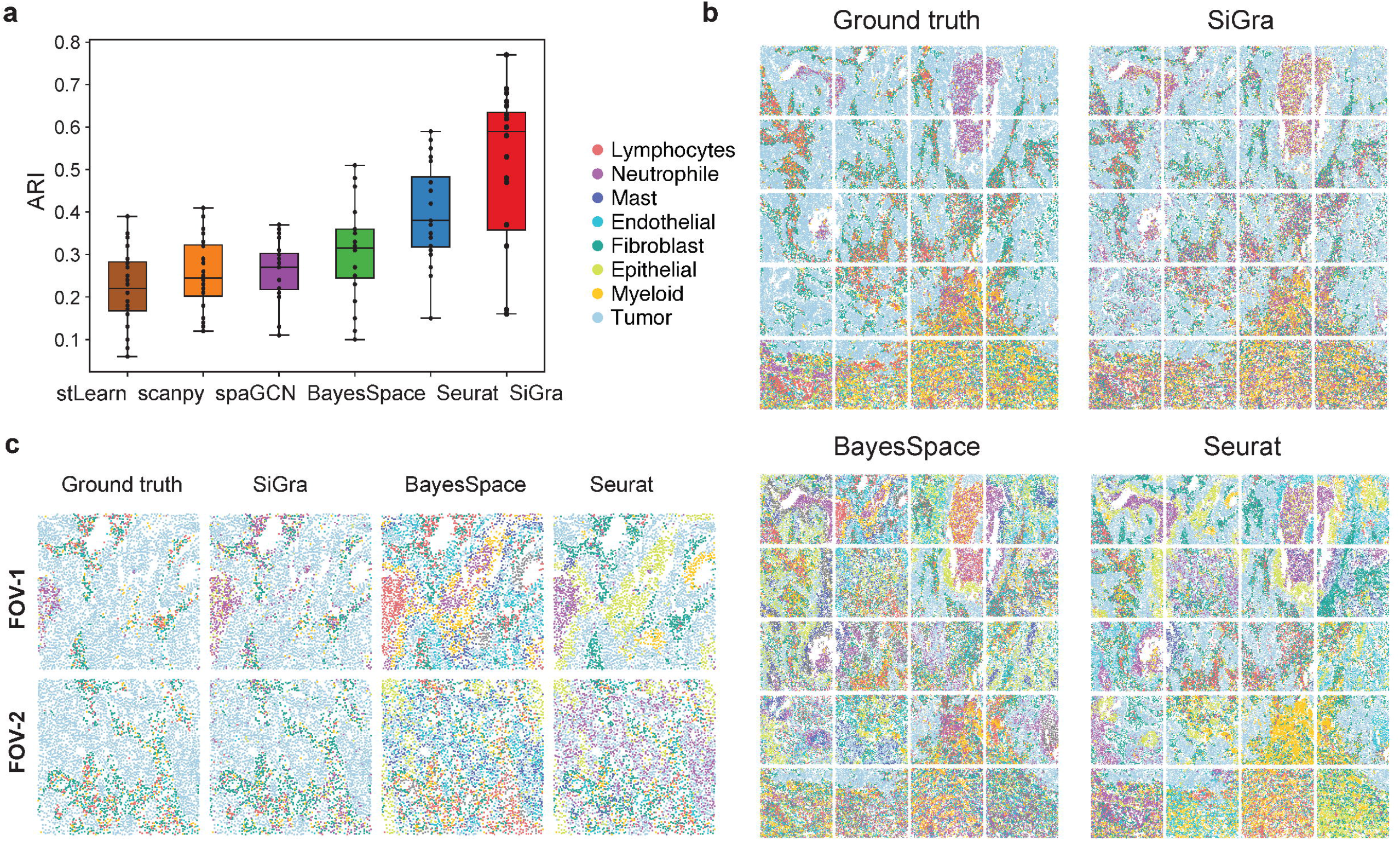
SiGra accurately identifies spatial domains in the single-cell spatial profiles of NanoString CosMx SMI. **a**, Boxplot of the adjusted rand index (ARI) scores of six methods in all the 20 FOVs of lung cancer tissue. The center line within the boxplot represents the median and the box limits denote the upper and lower quartiles. **b**, Spatial regions of ground truth and those detected by different methods including SiGra, BayesSpace, and Seurat. **c**, Spatial regions of two FOVs detected by different methods including SiGra, BayesSpace, and Seurat.

With the spatial clusters identified by different methods, the spatial organization of 20 FOVs are shown in **Fig. 2b**. Specifically, SiGra detects spatial domains that agree well with the original study, i.e., ground truth, with the overall ARI as 0.55, higher than Seurat (ARI = 0.37) and BayesSpace (ARI = 0.23). Seurat and BayesSpace significantly mislabel more cells than SiGra across the 20 FOVs. Meanwhile, the other two methods show much lower accuracy (ARIs: 0.25 for Scanpy, 0.22 for SpaGCN, and 0.34 for stLearn). Of note, the addition of multi-channel images (ARI = 0.59) improves the performance by 47.5%, compared with using gene expression only as input (median ARI = 0.40, **Supplementary Fig. 2a**). These results demonstrate that multi-modal spatial information contributes to the superior performance of SiGra.

The spatial clustering results are further scrutinized at FOV level (**Fig. 2c**). Of note, SiGra shows consistency between its identified cellular anatomy and the ground truth, with the continuous tumor region infiltrated with scattered immune cell clusters. In contrast, BayesSpace and Seurat misidentify the cellular anatomies as either a mixture of fragmental cell regions (FOV-1) or with highly blended cell types (FOV-2). For FOV-1, BayesSpace misidentifies the neutrophile as lymphocytes, meanwhile, it incorrectly identifies some tumor cells as myeloid cells or neutrophile. Seurat fails to disentangle epithelial cells from tumor cells. In FOV-2, BayesSpace misrecognizes fibroblast as the mixture of myeloid cells and lymphocytes, while Seurat mixes neutrophile with tumor cells without clear dissection of spatial heterogeneity. Those results indicate that the compared methods lack the capability of deciphering major spatial regions in single-cell spatial transcriptomics data.

### SiGra enhances gene expression patterns that distinguish intratumoral spatial heterogeneity

SiGra enhances the spatial gene expression data and improves downstream analysis for unveiling biological relevance. Herein, we perform the Uniform Manifold Approximation and Projection (UMAP) on raw data and enhanced data respectively (**Fig. 3a, Supplementary Fig. 2b**). Apparently, enhanced data reveals better data topology with different cell types better separated in the UMAP. Moreover, the enhanced cell type specific gene markers show prevalently consistent expression in their corresponding cell types (**Fig. 3b**). For example, the enhanced fibroblast marker gene *DCN*^18^ demonstrates uniform high expression in fibroblasts and low expression in other cell types. In contrast, in raw data, *DCN* presents sporadic expression in fibroblasts while highly expresses in other non-fibroblasts. Thus, SiGra not only denoises false-positive expressions (e.g. the *DCN* expression in non-fibroblasts) and extreme values, but also imputes missing values (e.g. the missing values of *DCN* expression in fibroblasts). Meanwhile, SiGra-enhanced data exhibits topological expressions of cell type specific markers (**Fig. 3c**). For example, after enhancement, *CD68*^19^ and *MGP*^20^ show elevated expressions in myeloid and endothelial cell enriched regions, respectively. Tumor-specific genes *EPCAM*^21^, *SOX4*^22^, and *KRT7*^23^, show strong and uniform enrichment in tumor regions, while these genes are not captured in the raw data of some tumor cells (**Fig. 3d**). In addition to marker genes, SiGra also allows revealing biologically meaningful differentially expressed genes (DEGs) (**Supplementary Fig. 2c**). Collectively, these results demonstrate the capability of SiGra to enhance gene expression data for better spatial data characterization.

**Fig. 3:**
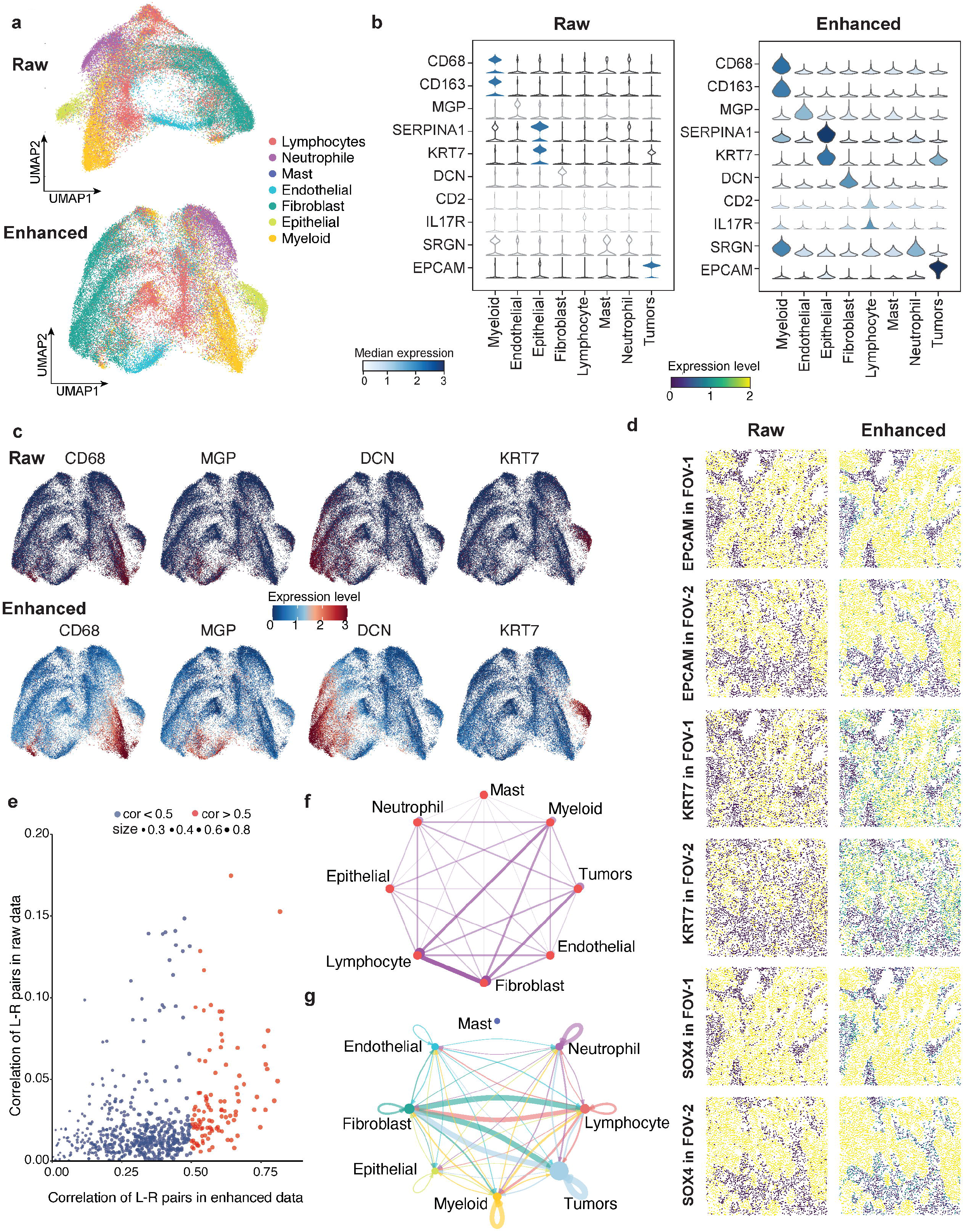
SiGra enhances gene expression patterns that distinguish intratumoral spatial heterogeneity. **a**, UMAP visualizations of raw data and the enhanced data of non-tumor cell population. **b**, Violin plots of the raw expression and the enhanced ones of marker genes in different cell populations. **c**, UMAP visualization of the raw expressions and the enhanced ones of marker genes (*CD68, MGP, DCN, KRT7*). **d**, Spatial visualization of the raw expressions and the enhanced ones of tumor related genes (*EPCAM, KRT7, SOX4*) in two FOVs. **e**, Scatter plot of the associations of all detected ligand-receptor (L-R) pairs in the raw data and the enhanced data, based on the non-tumor cell population. Red colored dots represent the L-R pairs that show strong association in the enhanced data. All L-R pairs were not identified with associations based on the raw data. **f**, Network representation of the weighted adjacent cell types. The edge width represents the adjacency between two cell types. **g**, Adjacent cell communications considering both neighboring cell weights and ligand-receptor interaction strength.

To further prove that the enhanced data by SiGra is useful for downstream analysis, we collect a combined list of candidate ligand-receptor (L-R) pairs from IUPHAR (International Union of Pharmacology)^24^, Connectome^25^, FANTOM5^26^, HPRD^27^, and Human Plasma Membrane Receptome (HPMR)^28^, and Database of Ligand-Receptor Partners (DLRP)^29^, which encompasses 815 ligands, 780 receptors, and 3,398 reliable L-R interaction pairs. Among them, genes of 660 L-R pairs are included in this NanoString CosMx dataset. For each of the 660 L-R pairs, we calculate the association of the ligand and its corresponding receptor, using raw data and enhanced data respectively. As shown in **Supplementary Fig. 3a**, of note, 221 L-R pairs with strong correlations (Pearson correlation > 0.5) are identified from the enhanced data. Surprisingly, no L-R pairs are observed with strong correlations based on raw data. Meanwhile, in the non-tumor cell population, we observe 104 L-R pairs with strong associations (**Fig. 3e**), where the classic *EFNB2* - *PECAM1* interaction^30^ is shown as one top associated L-R pair (Pearson correlation = 0.78, **Supplementary Fig. 3b**) in enhanced data. Both *EFNB2* and *PECAM1* present extensive zeros in raw data, with only a small portion of expressed cells observed, which explains the low correlation (Pearson correlation = 0.08) of *EFNB2* - *PECAM1* in raw data. Thus, the enhanced data co-expression patterns of L-R pairs have great biological relevance that facilitate the mining the cellular communications. Specifically, we further reveal the adjacency between different cell types (**Fig. 3f, Materials and Methods)** as well as the cell-cell communications considering both the neighboring cell weights and the L-R communication strength (**Fig. 3g**). Tumor-associated fibroblasts play a central role in the tumor microenvironment, not only adjacent to tumor cells and lymphocytes (**Fig. 3f**), but also presenting strong communications with them (**Fig. 3g**). In contrast, lymphocytes and myeloid cells are close with each other but with less communications. Collectively, this evidence demonstrate that the enhanced data contributes to downstream analysis and are crucial for revealing cellular interactions, which otherwise will be hidden due to data sparsity.

### SiGra enhances the single-cell spatial data of Vizgen MERSCOPE

SiGra is further evaluated on the other SCST dataset of mouse liver profiled by Vizgen MERSCOPE, which consists of 347 genes and 395,215 cells. In this dataset, SiGra reveals different spatial cell clusters (**Supplementary Fig. 4a**). For better visualization, we focus on the four major cell clusters (**Fig. 4a**). Cluster 1 (C-1) and cluster 2 (C-2) are located adjacent to central and portal veins respectively, while cluster 3 (C-3) and cluster 4 (C-4) are co-located with blood vessels. Importantly, the enhanced data by SiGra reveals histologically meaningful liver-specific gene expression patterns in different regions (**Fig. 4b**). For example, SiGra realizes remarkable enhancement of hepatocyte’s hallmark genes *Cyp2c38*^31^ and *Axin2*^32^, which are predominantly expressed near blood vessels. The endothelial cell markers *Cd34*^33^ (**Fig. 4b**) and *Vwf*^34^ (**Supplementary Fig. 4b**) also clearly present at central veins, portal veins, and sinusoids. The raw data, in contrast, shows noisy expressions of these genes in the non-relevant anatomic regions. For example, *Cd34* and *Vwf* show scattered false signals in the non-blood-vessel regions and missing expressions in smaller veins especially sinusoids. Thus, essential cellular anatomic structures in the liver tissue, such as central veins, portal veins, and sinusoids, can be clearly identified by the enhanced expressions of *Cd34, Vwf*, and *Axin2*, but not the noisy raw data, which are further confirmed by the UMAP plots (**Fig. 4c**). From the boxplots of these hallmark gene expressions (**Fig. 4d**), both C-1 and C-2 are suggested to be hepatocytes (high *Cyp2c38* and *Axin2* expressions), C-2 is enriched with periportal hepatocytes (higher *Axin2* expression), C-4 contains mainly endothelial cells (high *Cd34* and *Vwf* expressions), and C-3 is likely to be hepatic stellate cells. Notably, the enhanced cell-type specific genes only enrich in their restricted regions but not in irrelevant regions, suggesting that SiGra does not introduce noticeable artifacts in the enhanced data.

**Fig 4:**
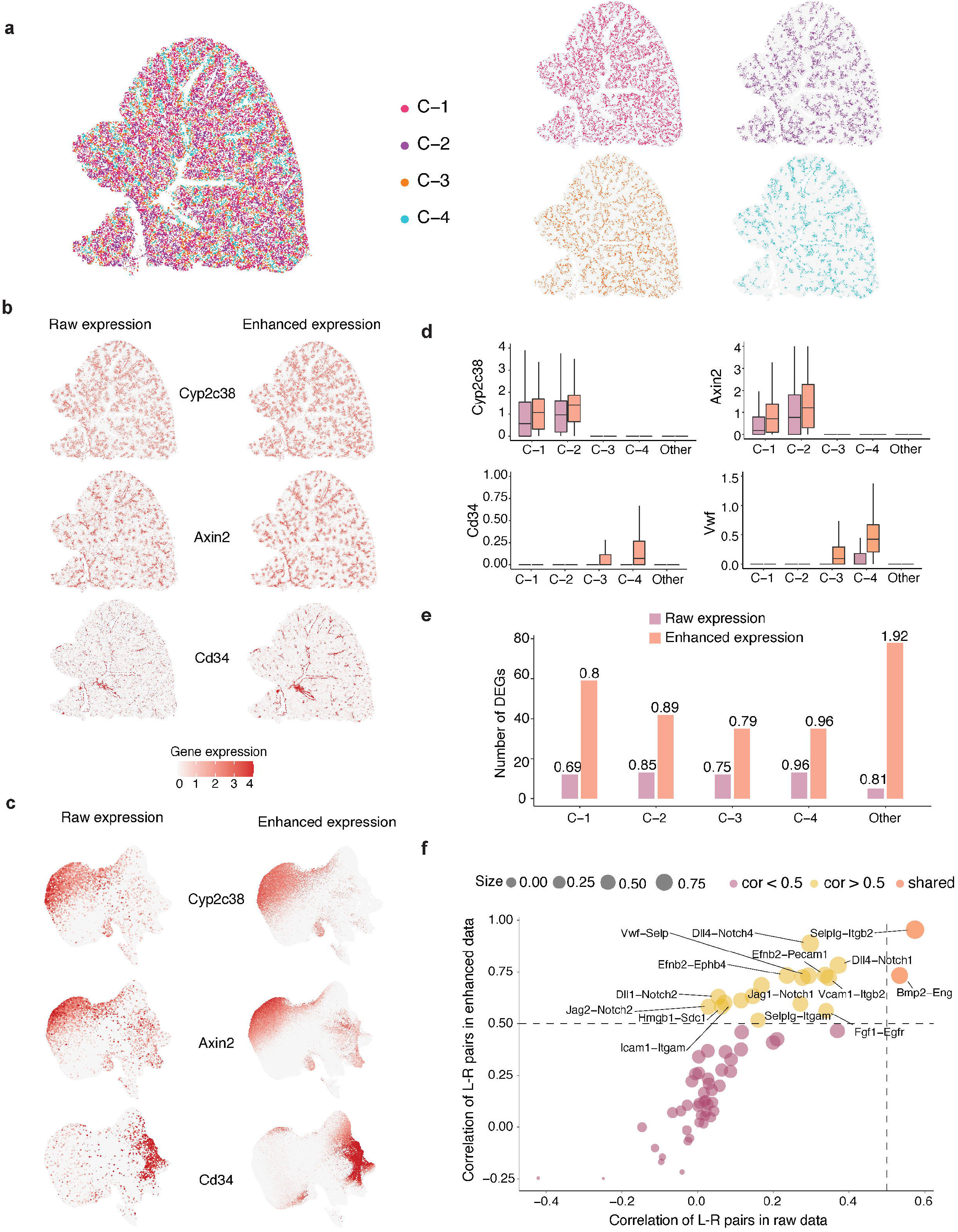
SiGra enhances the single-cell spatial data of Vizgen MERSCOPE. **a**, Spatial visualization of the cell clusters in the single-cell spatial data from mouse liver tissue. **b**, Spatial visualization of the raw expressions and the enhanced ones of liver related genes (*Cyp2c38, Axin2, Cd34*). **c**, UMAP visualization of the raw expressions and the enhanced ones of liver related genes (*Cyp2c38, Axin2, Cd34*). **d**, Boxplots of the raw expression and the enhanced ones of liver specific genes in different cell clusters. **e**, Comparisons of the number of differentially expressed genes (DEG) in cell clusters. The labeled number is the average log2FC for that cell cluster. **f**, Scatter plot of the associations of detected ligand-receptor (L-R) pairs in the raw data and the enhanced data. Yellow colored dots represent the L-R pairs that show strong association only in enhanced data.

In addition, SiGra reveals more differentially expressed genes (DEGs) from the enhanced data than raw data (**Fig. 4e**). For example, the enhanced data recovers 59, 42, 35 DEGs for C-1, C-2, and C-3, while the raw data only identifies 12, 13, 12 DEGs accordingly. The identified DEGs in enhanced data also show higher average log2 Fold Change (logFC; C-1: 0.8; C-2: 0.89; C-3: 0.79) than raw data. Of note, the enhanced data by SiGra reveals the associated ligand-receptor (L-R) pairs. As shown in **Fig. 4f**, among the 64 L-R pairs identified in this dataset, 19 pairs present strong correlations (Pearson cor > 0.5) based on the enhanced data. In contrast, only 2 L-R pairs are observed with strong correlations in raw data. *Selplg* (ligand) - *Itgb2* (receptor) is shown as the top associated L-R pairs both in enhanced data and raw data, while *Dll4* (ligand) - *Notch4* (receptor) presents strong association only in enhanced data. This result demonstrates that SiGra enhances gene expressions that facilitates recovering the liver-specific genes and potential cell-cell interactions, which enables better characterization of the spatial architecture of mouse liver tissue.

### SiGra improves identification of known layers in brain tissues

To show that SiGra not only outperforms existing methods in single-cell spatial data, but also in spot-based spatial transcriptomics data, here we analyze the 10x Visium datasets from human dorsolateral prefrontal cortex (DLPFC). These datasets consist of 12 tissue slices of human brains, covering up to six neuronal layers and white matter manually annotated by the original study. To evaluate the benchmarking performance, the identified spatial clusters are annotated based on the matched overlap of spatial clusters and ground truth. **Fig. 5a** shows the ARI scores for all 12 tissue slices, of which SiGra (median ARI: 0.54) outperforms Scanpy (median ARI: 0.28), Seurat (median ARI: 0.29), stLearn (median ARI: 0.39), SpaGCN (median ARI: 0.40), and BayesSpace (median ARI: 0.44).

**Fig. 5:**
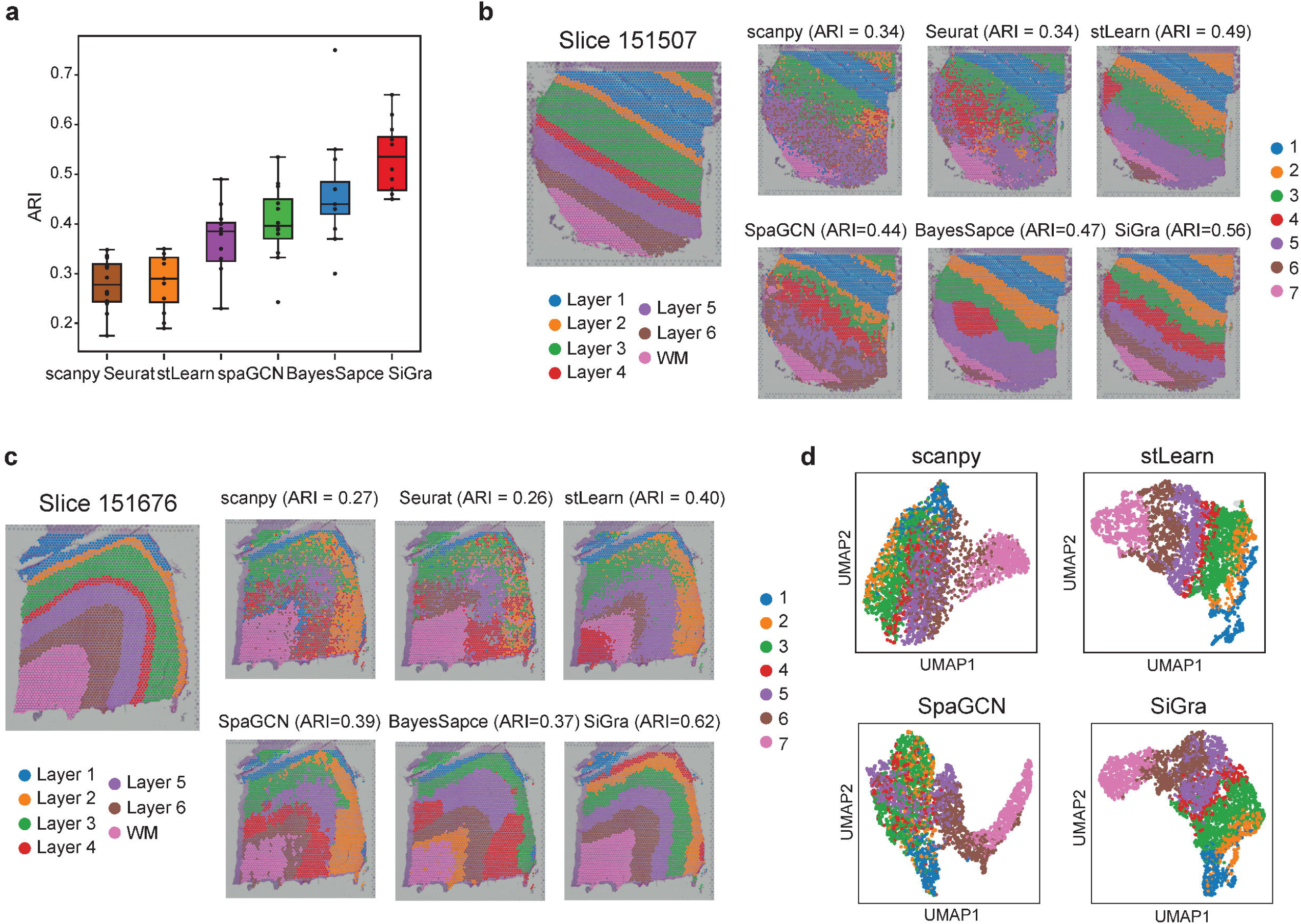
SiGra improves identification of known layers in brain tissues. **a**, Boxplot of the adjusted rand index (ARI) scores of six methods in all the 12 DLPFC slices. The center line within the boxplot represents the median and the box limits denote the upper and lower quartiles. **b**, Spatial domains detected by different methods including Scanpy, Seurat, stLearn, BayesSpace, SpaGCN, and SiGra, in the DLPFC slice 151507. **c**, Spatial domains detected by different methods including Scanpy, Seurat, stLearn, BayesSpace, SpaGCN, and SiGra, in the DLPFC slice 151676. **d**, UMAP visualizations of latent embeddings generated by Scanpy, stLearn, SpaGCN, and SiGra in the DLPFC slice 151676. SpaGCN and BayesSpace were not shown as they didn’t provide latent embeddings for UMAP visualization.

We further examine the DLPFC anatomic structures identified by different methods. For tissue slice 151507 (**Fig. 5b**), SiGra reveals more accurate spatial regions than other methods. Seurat identifies Layer 4 scattered in the regions of Layer 3 and 5 without clear boundaries. Scanpy, stLearn, and BayesSpace are not able to distinguish the anatomic shape of Layer 4. **Fig. 5c** shows the other tissue slice 151676 with spatial regions identified by different methods. Only SiGra deciphers the layer boundaries clearly that reaches good agreement with manual annotations (ARI = 0.62), while other methods can only achieve ARI less than 0.4. Specifically, stLearn intermingles Layer 2 with Layer 3, with additional mixtures of Layer 4 and white matter. BayesSpace mixes Layer 4 with Layer 5, and misidentifies some white matter as Layer 2, which leads to its poor performance. Interestingly, though stLearn also utilizes histology information from the H&E images to capture morphological features, its performance is substantially worse than SiGra, suggesting that SiGra incorporates the multi-modal spatial features in a more effective way. In addition, based on the latent embeddings of slice 151676 obtained by different methods (**Fig. 5d**), SiGra presents much clearer separations of different anatomic layers, while Scanpy and SpaGCN only discern white matters, failing to distinguish other neuronal layers. All these benchmarking results show that SiGra is able to better identify subtle spatial domains than other methods on the spot-based spatial transcriptomics data.

### SiGra improves spatial gene expression for better structural characterization

To validate further that SiGra enhances the spatial gene expressions, we detect the DEGs of each domain in slice 151676. Compared with the number of DEGs detected in raw data, SiGra detects more DEGs specific to individual regions (**Supplementary Table 1**). For example, with 232 DEGs of Layer 1 detected in raw data, 595 DEGs are found specific to Layer 1 from enhanced data. Moreover, for each of the neuronal layers, region-specific maker genes can be better identified after SiGra data enhancement (**Fig. 6a**). For example, *MYH11*^35^ presents enriched expressions in Layer 1 (logFC = 2.96). *C1QL2*^36^ and *CUX2*^36^ are overexpressed in Layer 2 and Layer 3, with logFC as 1.74 and 1.44. *SYT2*^37^ and *FEZF2*^38^ are enriched in Layer 4 (logFC = 1.31) and Layer 5 (logFC = 1.42). *PAQR6*^39^ shows dominantly enriched expressions (logFC = 2.9) in white matter area. In contrast, these markers genes do not show clear expression patterns in raw data, indicating the limits that raw data faces in distinguishing spatial domain boundaries. Violin plots further show the expressions of marker genes in raw data and enhanced data respectively (**Fig. 6b**). Such enhanced gene expression patterns are also observed in other DLPFC slices, for example, slice 151507 (**Fig. 6c**), where *RELN*^40^ (logFC = 3.24) and *ADCYAP1*^41^ (logFC=2.81), i.e. makers of Layer 1 and Layer 2, present remarkable enhancement, in contrast to their sporadic expressions in raw data. The enhanced data by SiGra consistently reveals more DEGs compared to the raw data (**Supplementary Table 2**). These results demonstrate the capability of SiGra to reduce noises and improve gene expression patterns in spot-based spatial transcriptomics data.

**Fig. 6:**
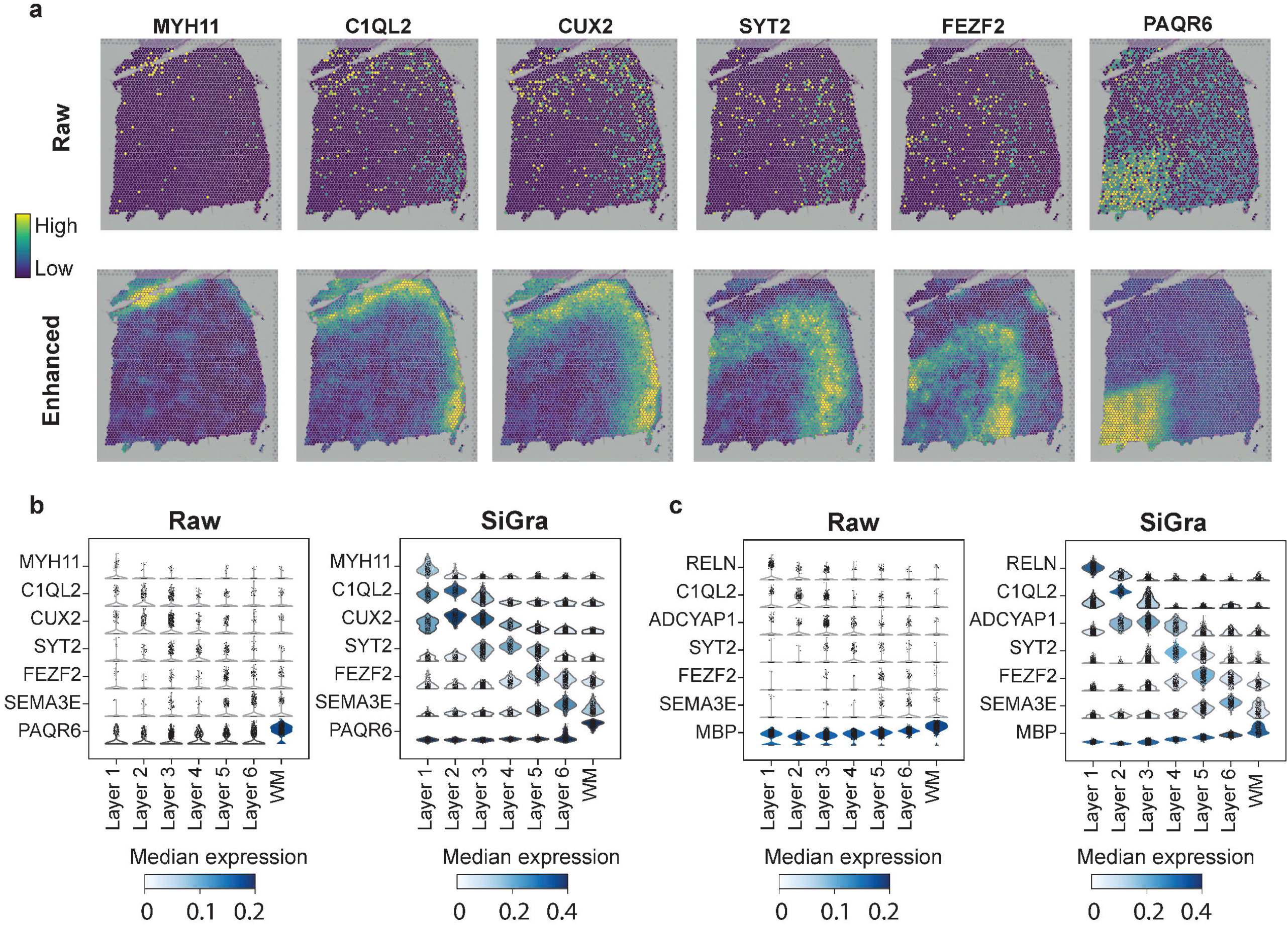
SiGra improves spatial gene expression for better structural characterization. **a**, Spatial visualization of the raw expressions and the enhanced ones of marker genes (*MYH11, C1QL2, CUX2, SYT2, FEZF2, PAQR6*) in DLPFC slice 151676. **b**, Violin plots of the raw expression and the enhanced ones of marker genes in DLPFC slice 151676. **c**, Violin plots of the raw expression and the enhanced ones of marker genes in DLPFC slice 151507.

## DISCUSSION

The recent spatial biology technologies have rapidly evolved into the single cell era^42^. Commercially available in situ hybridization platforms such as NanoString CosMx SMI and Vizgen MERSCOPE have enabled spatial gene expression profiling at subcellular resolution (50nm) for 500 – 1,000 targeted genes. Experimental in situ sequencing technologies such as ExSeq^43^ expand SCST to transcriptome-wide. Spot-array spatial transcriptomics technologies such as Stereo-seq^44^ and Seq-Scope^45^ are also reaching subcellular resolution (500nm – 600nm). However, common in all these technologies, the resulting SCST data is limited by the low total transcripts per cell, noisy data, and substantial zeros, which raises challenges in effective downstream analysis^42^. To accurately reveal spatial and cellular anatomic structures and to enhance the noisy gene expression data, we have developed the SiGra method, a graph artificial intelligence model, to incorporate multi-modal data including multi-channel IHC images, spatial adjacent cell graph, and gene expressions. The use of the graph transformers over the spatial adjacent cell graph as well as the imaging-transcriptomics hybrid architecture allow SiGra to effectively leverage the rich information from the high-content IHC images as well as the spatial distribution. In SiGra, the multi-modal information from images and original transcriptomics are summarized at single-cell level, with the information from neighboring cells selectively captured by the attention mechanism. With these technical advances, the SiGra model outperforms existing methods and significantly improves downstream data analysis.

SCST grants researchers the spatial perspective for exploring the cellular ecosystems in complex tissues. SiGra is one of the first method to utilize multi-modalities including multi-channel images of cell morphology and functions to address technology limitations and achieve augmented spatial profiles. Coupling the unique transcriptomics profiles with multi-channel imaging data improves the interpretability of the spatial transcriptomics data and gains molecular-level insights in cellular pathology.

SiGra is designed as a general-purpose tool for spatial transcriptomics data enhancement and spatial pattern profiling. Besides the SCST data, SiGra can also be directly used for spot-based spatial transcriptomics data such as the 10x Visium and demonstrates superior performance than existing methods. SiGra provides platform-specific preprocessing tools for CosMx, MERSCOPE, and 10x Visium data. The enhanced SCST data by SiGra are ready-to-use for downstream bioinformatics analyses. SiGra has demonstrated superior performance over three different platforms, in both health and disease tissues, and across different species. Therefore, SiGra provides a general solution for existing spatial transcriptomics data analysis pipelines.

Besides the superior performance and technical advantages, SiGra can be further improved in the future. First, as newer spatial omics technologies^46^ continue evolving and new data modalities keep emerging, SiGra can be improved by incorporating new omics data types, new image types, 3-D spatial information, etc., to extend the data exploration. The hybrid architecture allows SiGra to adapt additional spatial information and incorporate multi-omics data. As a advanced deep learning model, SiGra also faces the limitations of the black-box nature of artificial intelligence^47-49^. This can be ameliorated through downstream analysis such as cell-cell interaction analysis. Further development of SiGra will enhance the model interpretability that can address some of the problems and bring insights into the underlying mechanism in tissue ecosystems. With the capacity and efficiency of the experimental technologies continue to improve, SiGra is anticipated to facilitate biological discoveries and insights into the complex tissues and diseases.

## MATERIALS AND METHODS

### Data Preprocessing and Graph representation

Spatial transcriptomics data generated by different platforms, including the NanoString CosMx^™^ SMI lung cancer dataset (Lung-9-1)^4^, Vizgen MERSCOPE mouse liver dataset L1R1 released in January 2022^15^, and 10x Visium datasets from human dorsolateral prefrontal cortex (DLPFC)^50^, are preprocessed and represented in the uniform format (**Fig. 1**) for SiGra. The original spatial profiles are converted to single-cell (or spot) images, single-cell (or spot) expression, and a spatial graph of adjacent cells (or spots), which serve as the input for SiGra.

Regarding the NanoString Lung-9-1 dataset, the composite images of the DAPI, PanCK, CD45, and CD3 channels from 20 FOVs, the cell center coordinates (from the cell metadata file), the single-cell gene expression file of 960 genes are used. For each cell, four images of 120-by-120 pixels (21.6-by-21.6μm) with the cell at the center are cropped from the images. The spatial adjacent graph is constructed based on the cell-to-cell distance (Euclidian distance) ≤ 80 pixels (14.4μm). NanoString’s annotations of cell types are obtained from their provided Giotto object. Regarding the Vizgen L1R1 dataset, images of the DAPI staining and the three IHC boundary staining, the single-cell expression data of 347 genes, and the cell center coordinates are used. The images in the middle of the z-packs (z3) are used, as recommended by Vizgen. These images are cropped into single-cell images with 200-by-200 pixels (21.6-by-21.6μm). The spatial adjacent graph is constructed with cell-to-cell distance ≤ 150 pixels (16.2μm). Regarding the 10x Visium DLPFC dataset, the high-resolution H&E images as well as the .h5 files (“filtered_feature_bc_matrix.h5”) are used as input. For each spot, three spot-specific images (for the RGB channels, respectively) are extracted, with the image size of 50-by-50 pixels (38.7-by-38.7μm). The cutoff distance for generating the spatial graph between spots is 150 pixels (116μm). Top 3000 highly variable genes are identified using Seurat standard pipeline^51^ and used for analysis. For all datasets, the raw counts of gene expressions are normalized by multiplying 10,000 and followed by log-transformation. The parameters of the size of single-cell images and the cutoff of cell-to-cell distance for constructing the spatial graph are determined empirically depending on cellular anatomy of the tissue. In the spatial adjacent graph, most cells have 5 – 6 neighboring cells.

In this way, the final graph representation of the original single-cell or spot spatial transcriptomics data is a spatial graph *G=*(*V,E*), with *v*_*i*_ ∈ *V*, representing the *i*’th cell with *i* = 1, …, *n* represent the totally *N* cells, *e*_*ij*_ ∈ *E* representing the spatial proximity between two cells *v*_*i*_ and *v*_*j*_, and *A* as the adjacent matrix of the graph. Each cell *v*_*i*_ on the spatial graph is accompanied with multi-channel images ***M***_*i*_ = {*M*_*i,c*_}, with *c* = 1, …, *C* representing each imaging channel, and gene expression ***g***_*i*_ = {*g*_*i,k*_}, with k = 1,…, *K* representing genes.

### The SiGra model

SiGra is a hybrid multi-modal graph transformer autoencoder with three transcriptomics reconstruction modules: the imaging-based autoencoder, the transcriptomics-based autoencoder, and the hybrid autoencoder.

1. Imaging-based autoencoder. For a cell *v*_i_, the multi-channel images ***M***_*i*_ are transformed to a vector ***x***_i_ ≡ [vec(*M*_i,1_), …, vec(*M*_*i,c*_)]^*T*^. An autoencoder with a series of multi-head graph transformer layers is used to project the imaging vector to the latent space as ***Z***_*M,i*_, then reconstruct the gene expression profile of this cell *v*_*i*_ as 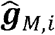. The images ***M***_*j*_ from neighboring cell *v*_*j*_ ∈ 𝒩(*v*_*i*_) are also used as the input, where 𝒩(·) represents the neighbors in the graph *G*.
2. Transcriptomics-based autoencoder. The original gene expression profile ***g***_*i*_ for cell *v*_*i*_ with ***g***_*i*_ for adjacent cells *v*_*j*_ ∈ 𝒩 (*v*_*i*_) also as the input, is projected to the latent space as ***Z***_*g,i*_, which is then used to reconstruct the gene expression of cell, which is then used to reconstruct the gene expression of cell *v*_*i*_ as 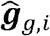.
3. Hybrid autoencoder. The latent representation of the imaging and the transcriptomics features are catenated as **[*Z***_*M,i*,_ ***Z***_*g,i*_ ], further projected to hybrid latent feature ***Z***_*h,i*,_ and then used to reconstruct the gene expression for cell *v*_*i*_ as 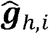 through graph transformer layers. The latent features ***Z***_*M,j*,_ and ***Z***_*g,j*_ from neighbor cells {*v*_*j*_} are also used by the graph transformers.

### Graph transformer convolutional layer

Multi-head graph transformer^52^ layers with attention mechanism (**Supplementary Fig. 1**) are the main components of the SiGra model. Briefly, the for a cell *v*_*i*_, the propagation of the graph transformer from the *l* layer to the *l* + 1 layer is defined as: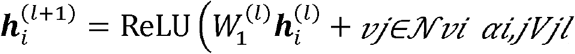, where the rectified linear unit (ReLU^53^) is used as the nonlinear gated activation function. The attention module is defined as: 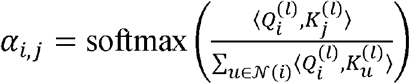, where:

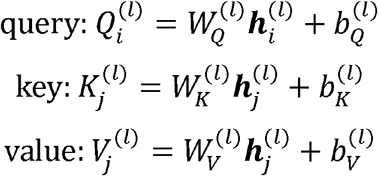

and 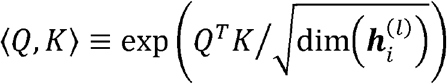. The multi-head attentions, which notation is omitted for simplicity, are concatenated.

### Loss function

SiGra learns the reconstructed gene expression via a self-supervised loss of combined MSE from gene embedding, image embedding and combined embedding, with the loss function

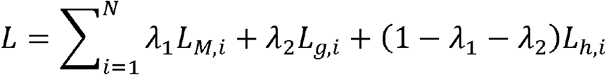

where 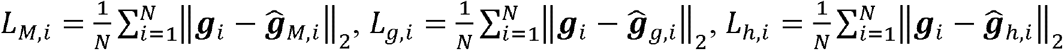. In this work, we set the hyperparameters *λ*_1_ = *λ*_2_ = 0.1.

The hyperparameters of SiGra are: two graph transformer layers for the imaging and the transcriptomics encoders (with dimension of 512 and 30 for the 1^st^ and the 2^nd^ layers, respectively), one graph transformer layer for the hybrid encoder (dimension of 60 with 30 from the transcriptomics encoder and 30 from the imaging encoder), and two graph transformer layers for imaging, transcriptomics, and hybrid decoders (with dimension of the first layer of 512, and the dimension of the second layer is same as the corresponding transcriptomics data). After training, SiGra outputs the hybrid reconstruction 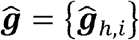 as the final enhanced expression profile. The latent representation, ***Z* =** { ***Z***_*h,i*_}, of the original SCST data is used for spatial data clustering with Leiden algorithm^54^ from the SCANPY package^11^.

### Benchmarking methods and comparison measurement

To evaluate the performance of SiGra, we compare it with five existing methods, including Seurat v4^10^, Scanpy^11^, stLearn^12^, SpaGCN^14^, and BayesSpace^13^. Seurat and Scanpy are implemented based on their provided vignettes. Briefly, for data preprocessing, 3,000 highly variable genes are selected for log normalization, and top 30 principal components (PCs) are calculated for spatial data clustering. BayesSpace is implemented based on their package vignette. Specifically, the input is the top 15 PCs of the log-normalized expression of the top 2,000 HVGs. The nrep parameter is set as 50,000 and the gamma parameter is set as 3. For stLearn, based on its tutorial, the stLearn.SME.SME_normalized() function is performed on raw counts with parameters use_data = “raw” and weights = “physical_distance”. The top 30 PCs of the SME normalized matrix are then used for spatial data clustering and visualization. SpaGCN is applied according to its recommended parameters in the package vignette. That is, the top 15 PCs of the log-normalized expression of the top 3,000 spatial variable genes are used for spatial data clustering. 200 epochs are used for identifying and refining spatial domains. The resolution parameter is selected to ensure the number of clustering is equal to the ground truth.

To evaluate the performance of each method, we use the Adjusted Rand Index (ARI) to assess the agreement between the identified spatial clusters and the manual annotation. Suppose 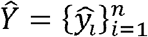 represent the spatial clusters and 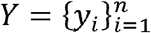 represent the ground truth of *n* cells with divided into *k* clusters. Then

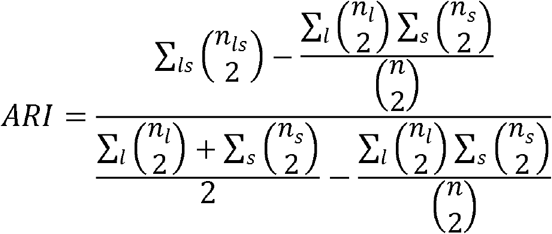

where *l* and *s* denote the *k* clusters, 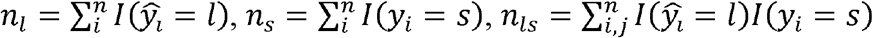, and *I* (*x* = *y*) = 1 when *x* = *y*, else *I* (*x* = *y*) = 0. The ARI ranges from 0 to 1 for increasing match between the identified clusters with ground truth.

### Identifying differentially expressed genes and adjacent cell communications

To identify the differentially expressed genes (DEGs), the Wilcoxon test from Scanpy package^11^ is used. DEGs of each spatial region is selected with 5% FDR threshold (Benjamin-Hochberg adjustment) and the log2 Folder Change more than 1 (log2 FC > 1).

To reveal the neighbors of each cell type, we aggregate its neighboring cells by cell type and divide the average number to reveal its weighted neighbors. These weighted neighbors are further used to characterize the adjacent cell communications. Specifically, the interaction strength of each L-R is calculated by the multiplication of their association score and their average expression. Then we aggregate the interaction strength of each L-R pair by cell type to be the communication strength of two cell types. The neighboring weight of two cell types is further multiplied to the communication strength of these two cell types for the final adjacent cell communications. In this way, the higher value of the adjacent communications, the stronger the two neighboring cell types interact.

## DATA AVAILABILITY

**NanoString CosMx SMI data:** This single-cell spatial dataset contains 20 FOVs, which is profiled by the CosMx SMI on Formalin-Fixed Paraffin-Embedded (FFPE) samples of the non-small-cell lung cancer (NSCLC) tissue^4^. The dataset (Lung-9-1) is available from https://nanostring.com/products/cosmx-spatial-molecular-imager/ffpe-dataset/. **Vizgen MERSCOPE/MERFISH dataset:** We used the Vizgen MERFISH Mouse Liver Map dataset that contains a MERFISH measurement of a 347 gene panel. Sample L1R1 (liver 1, replicate 1) was used and downloaded from https://info.vizgen.com/mouse-liver-access, which includes the list of detected transcripts, gene counts per cell matrix, additional spatial cell metadata, cell boundary polygons, and DAPI images. **10x Visium dataset of human brain:** The human dorsolateral prefrontal cortex (DLPFC) 10x Genomics Visium datasets^20^ consists of 12 samples. Each of the sample is manually annotated with up to six cortical layers and white matter. Transcriptomics data and hematoxylin and eosin (H&E) images of corresponding tissue sections are downloaded from http://research.libd.org/spatialLIBD/.

## CODE AVAILABILITY

SiGra is provided as a Python package available at https://github.com/QSong-github/SiGra, with detailed tutorials for the general applicability on different SCST platforms.

## AUTHOR CONTRIBUTIONS

QS, JS, and BY developed the structure and arguments and wrote the manuscript. ZT implemented the method and applications. All the authors reviewed and approved the final manuscript.

## MATERIALS & CORRESPONDENCE

Correspondence and requests for materials should be addressed to QS or JS.

## FIGURE LEGENDS

**Supplementary Fig. 1: Illustration of the multi-head graph transformer layer in SiGra. a**, the overall architecture of a graph transformer layer. **b**, the attention module in the graph transformer.

**Supplementary Fig. 2: Multi-modal based SiGra improves clustering accuracy and enhances spatial gene expression in lung cancer tissue. a**, Boxplot of the adjusted rand index (ARI) scores of SiGra based on image features and gene expression features in all the 20 FOVs of lung cancer tissue. The center line within the boxplot represents the median and the box limits denote the upper and lower quartiles. **b**, UMAP visualizations of raw data and the enhanced data of all cells. **c**, Violin plots of top differentially expressed genes in different cell populations.

**Supplementary Fig. 3: SiGra facilitates the discoveries of ligand-receptor interactions with biological significance. a**, Scatter plot of the associations of all detected ligand-receptor (L-R) pairs in the raw data and the enhanced data, based on all cell population. Red colored dots represent the L-R pairs that show strong association in the enhanced data. All L-R pairs were not identified with associations based on the raw data. **b**, Scatter plot of the raw expressions of top ligand-receptors, including *EFNB2* (ligand) - *PECAM1* (receptor) and *CD48* (ligand) - *CD2* (receptor) in raw data and enhanced data.

**Supplementary Fig. 4: SiGra reveals spatial cell distribution and enhances gene expression in mouse liver tissue. a**, Spatial visualization of all cell clusters in the single-cell spatial data from mouse liver tissue. **b**, Spatial visualization of the raw expressions and the enhanced expressions of *Vwf*.

**Supplementary Table 1. DEGs identified for each layer based on raw data and enhanced data of slice 151676**.

**Supplementary Table 2. DEGs identified for each layer based on raw data and enhanced data of slice 151507**.

